# Inhibition of HSP90 reversed STAT3 mediated muscle wasting induced by cancer cachexia

**DOI:** 10.1101/2021.01.27.428420

**Authors:** Mengyuan Niu, Shiyu Song, Zhonglan Su, Lulu Wei, Li Li, Wenyuan Pu, Chen Zhao, Yibing Ding, Wangsen Cao, Qian Gao, Hongwei Wang

**Author notes:** Address for Correspondence: Hongwei Wang, Center for Translational Medicine and Jiangsu Key Laboratory of Molecular Medicine, Medical School of Nanjing University, Nanjing 210093, PR China. Equal contribution (Mengyuan Niu, Shiyu Song).

## Abstract

Cancer cachexia is one of the most common causes of death among cancer patients, no effective anti-cachectic treatment is currently available. In experimental cachectic models, aberrant activation of STAT3 in skeletal muscle has been found to contribute to muscle wasting. However, its clinical association, the factors regulating STAT3 activation, and the molecular mechanisms of STAT3-induced muscle atrophy in cancer cachexia remain incompletely understood. Here, we show that an enhanced interaction between STAT3 and HSP90, which causes the persistent STAT3 activation in the skeletal muscle of cancer cachexia patients, is the crucial event for the development of cachectic muscle wasting. Administration of HSP90 inhibitors alleviated the muscle wasting in C26 tumor-bearing cachetic mice model or C26 conditional medium induced C2C12 myotube atrophy. A mechanistic study indicated that in cachectic skeletal muscle, prolonged STAT3 activation triggered muscle wasting in a FOXO1-dependent manner, STAT3 activated FOXO1 by binding directly to its promoter. Our results provide key insights into the role of the HSP90/STAT3/FOXO1 axis in cachectic muscle wasting, which shows promising therapeutic potential as a target for the treatment of cancer cachexia.

## Introduction

Cancer cachexia is a devastating metabolic syndrome that affects up to 80% of cancer patients and accounts for nearly 20% of all cancer-related deaths (Loberg *et al*, 2007). Skeletal muscle wasting is one of the most crucial pathological events in cancer cachexia. As a highly plastic tissue, skeletal muscle can change its mass, function, and metabolism in response to various endogenous or exogenous stimuli. Current evidence indicates that the imbalance between catabolic and anabolic responses and the disorder of protein synthesis and degradation pathways, including both the ubiquitin-proteasome system (UPS) and the autophagy-lysosome pathway (ALP), are the major causes of cachectic muscle wasting (Bilodeau *et al*, 2016). Activation of the UPS is a crucial event in muscle wasting in cancer cachexia, and the enhanced expression of muscle-specific E3 ubiquitin ligases, such as muscle-atrophy-F-box (MAFbx/atrogin-1) and muscle-RING-finger-1 (MuRF1), are hallmarks of this process (Bodine *et al*, 2001; Guadagnin *et al*, 2018; Guo *et al*, 2017). However, the upstream activators of the protein degradation pathway and the molecular mechanisms involved in muscle wasting in cancer cachexia are still largely unknown.

Signal transducer and activator of transcription 3 (STAT3) plays a critical role in cancer cachexia, and increased STAT3 activation (in muscle) has been found in multiple types of experimental cancer cachexia (Zimmers *et al*, 2016)(Ma *et al*, 2017; Mubaid *et al*, 2019; Silva *et al*, 2015). In experimental cancer cachexia, muscle-specific STAT3 depletion or JAK2/STAT3 pathway inhibition can reverse the skeletal muscle wasting phenotype (Bonetto *et al*, 2012). However, its clinical association, the reason for prolonged STAT3 activation in cachectic muscles, and its contribution to pathological anabolic responses in muscle still need to be defined.

HSP90 (heat shock protein 90) is an evolutionarily conserved molecular chaperone that is essential for cell growth, proliferation, transformation, proliferation, and survival under normal and stress conditions (Schopf *et al*, 2017). HSP90 interacts extensively with a variety of signaling transduction proteins (Whitesell *et al*, 2005),(Mahendrarajah *et al*, 2017). We and other groups have previously demonstrated that HSP90 could directly interact and regulate STAT3 activation in various cancers and promote cancer cell growth and survival as an oncogene (Prinsloo et al. 2012; Schoof et al. 2009; Song et al. 2017b). However, whether HSP90 is involved in regulating STAT3 activation in skeletal muscles and its functional role in cachectic muscle wasting is still unknown.

Herein, we report that the increased HSP90-STAT3 interaction is necessary to induce prolonged STAT3 activation and muscle atrophy in clinical cachetic patients and C26 tumor-bearing experimental cachexia mice model. Administration of HSP90 inhibitors *in vivo* could successfully alleviate the pathological development of experimental cachexia; mechanistic study demonstrated that activated STAT3 could induce FOXO1 transcription by binding directly to the FOXO1 promoter, and knockdown of FOXO1 abolished the effects of STAT3 induced muscle wasting. Therefore, our study provided novel experimental evidence showing that the HSP90/STAT3/FOXO1 axis might be a potential therapeutic target for cancer cachexia.

## Results

### Enhanced HSP90-STAT3 interaction causes the persistent STAT3 activation and muscle atrophy in patients and mice with cancer cachexia

We first evaluated the expression level of HSP90, atrophy markers, and activation of STAT3 in muscle from cancer cachexia patients. As shown in Fig 1, there is a significantly increased expression of muscle atrophy markers, including myostatin, E3 ligases such as Atrogin-1 and MuRF1 in skeletal muscle of cancer cachexia patients; also we detected the persistent activation of STAT3 accompanied by significantly increased expression of HSP90 in cachetic skeletal muscle (Figure 1A, B, C); by measuring the HSP90-STAT3 interaction using the immunoprecipitation, we observed a dramatic increase in the HSP90-STAT3 interaction in patient muscle with cancer cachexia (Figure 1D), indicating that HSP90 might be involved in the regulation of STAT3 activation during the pathological development of cancer cachexia.

**Figure 1.**
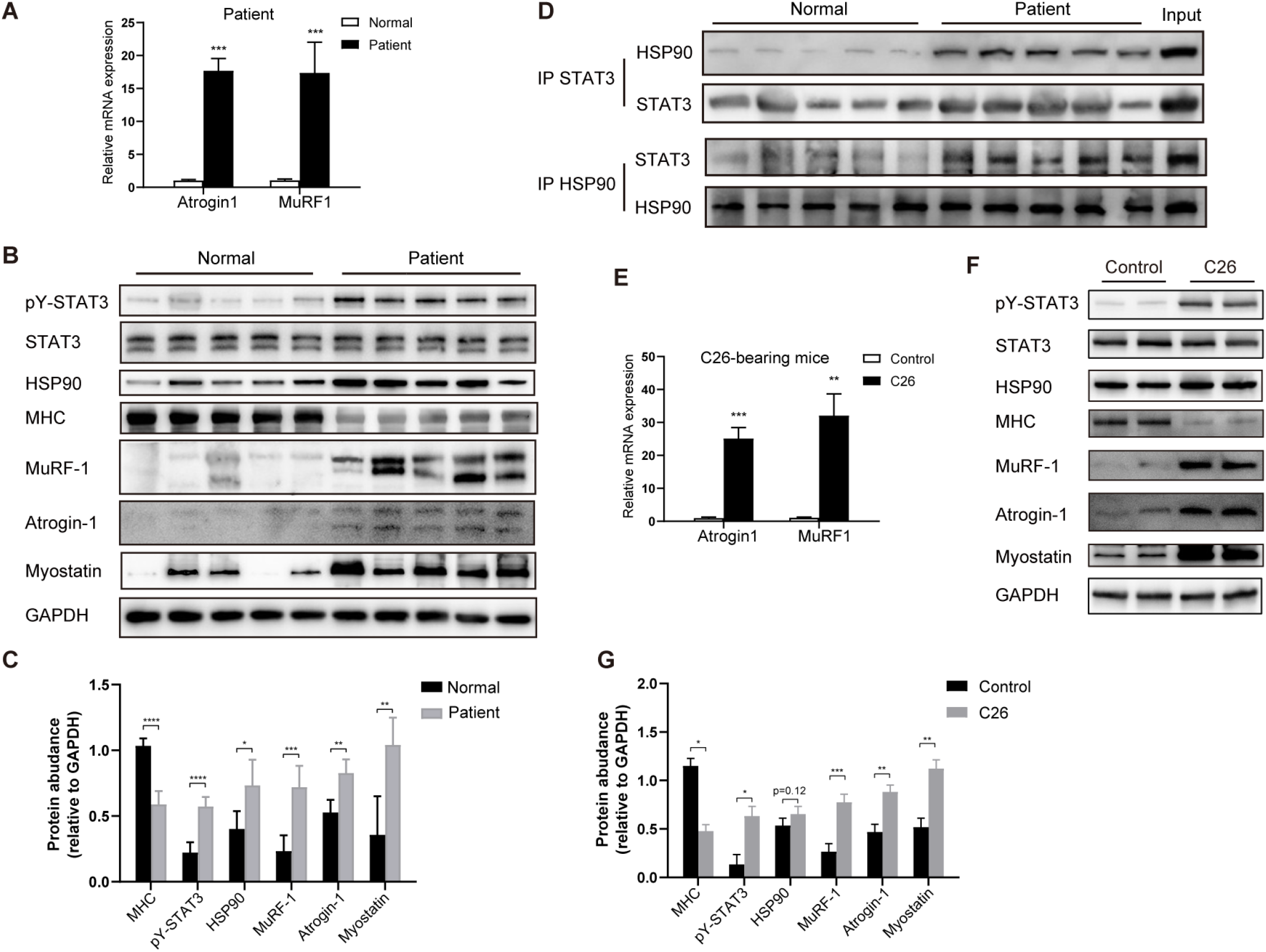
Enhanced HSP90-STAT3 interaction induced prolonged STAT3 activation were detected in the skeletal muscle from cancer cachetic patient and C26 tumor-bearing cachetic mice. The expression levels of Atrogin-1, MuRF-1, MHC, HSP90, Myostatin, and phosphorylated STAT3 in the skeletal muscle from cancer cachetic patients were detected by qRT-PCR (A. For atrogin-1 and MuRF-1) and Western blot (B), the bands were quantified by densitometry and normalized to the density of GAPDH (C). Measurement of the HSP90-STAT3 interaction in cachetic muscle cancer cachetic patients by immunoprecipitation (D). The expression levels of Atrogin-1, MuRF-1, MHC, HSP90, Myostatin and phosphorylated STAT3 in the skeletal muscle from C26 tumor-bearing cachectic mice were detected by qRT-PCR (E. For atrogin-1 and MuRF-1) and Western blot (F), the bands were quantified by densitometry and normalized to the density of GAPDH (G).

To validate these results, an experimental cachexia mouse model was adopted by inoculated Balb/C mice with a mouse colon adenocarcinoma cell line C26; these mice showed typical cachectic symptoms, including progressive weight loss, dull and matted fur, and mental weakness, which was consistent with the previous reports (Tanaka *et al*, 1990). There was approximately 15-20% overall weight loss and obvious lean body mass weight loss in C26 tumor-bearing mice compared with that in non-tumor-bearing mice within 15 days after tumor inoculation (Fig EV1A). Persistent STAT3 activation, as well as the enhanced expression of HSP90 and atrophy related molecular markers including myostatin, Atrogin-1, and MuRF1, were also detected in cachectic muscle from C26-bearing mice (Fig 1E, F, G), the dramatic increase in the HSP90-STAT3 interaction were also confirmed in the cachetic muscle in C26 tumor-bearing mice (Fig EV1B).

### HSP90 and STAT3 were required for C26 CM-induced myotube atrophy

To assess the involvement of HSP90 in cancer cachexia-related muscle wasting, we adopted the *in vitro* cachectic cell model by culturing the C2C12 myotubes with C26 tumor cell-derived conditional medium (C26 CM)(Fig EV1C). Western blot assay confirms the pathological phenotype of cachetic muscle atrophy based on the loss of MHC expression and the enhanced expression of myostatin, E3 ligases such as Atrogin-1 and MuRF1 (Fig 2A, 2B). H&E staining further demonstrated the myotube shrinking induced by C26 CM (Fig 2D). Notably, we observed a dramatic increase in the HSP90-STAT3 interaction in C2C12 myotubes upon C26 CM treatment (Fig 2C); this is consistent with the *in vivo* data, although the increased expression of HSP90 in skeletal muscle was not obvious in C26 CM induced muscle atrophy (Fig 2A). To determine whether the atrophy effect was HSP90-dependent, we knocked down HSP90 in C2C12 myotubes. The results showed that the administration of RNAi against HSP90 or STAT3 prevented the decrease in the myotube size, myotube diameter and myotube area induced by C26 CM (Fig 2D,E).These results were further confirmed by the measurement of the transcriptional expression of MuRF-1 and atrogin-1 (Fig 2E right). Consistently, the myotube atrophy-associated phenotypes, including the downregulated expression of MHC and the upregulated expression of myostatin, MuRF-1, and atrogin-1, were largely reversed when HSP90 was knocked down (Fig 2F left). Similar effects were observed when STAT3 was knocked down in C2C12 myotubes (Fig 2F right), indicating that the enhanced HSP90-STAT3 interaction plays a crucial role in the pathological development of cachetic muscle wasting.

**Figure 2.**
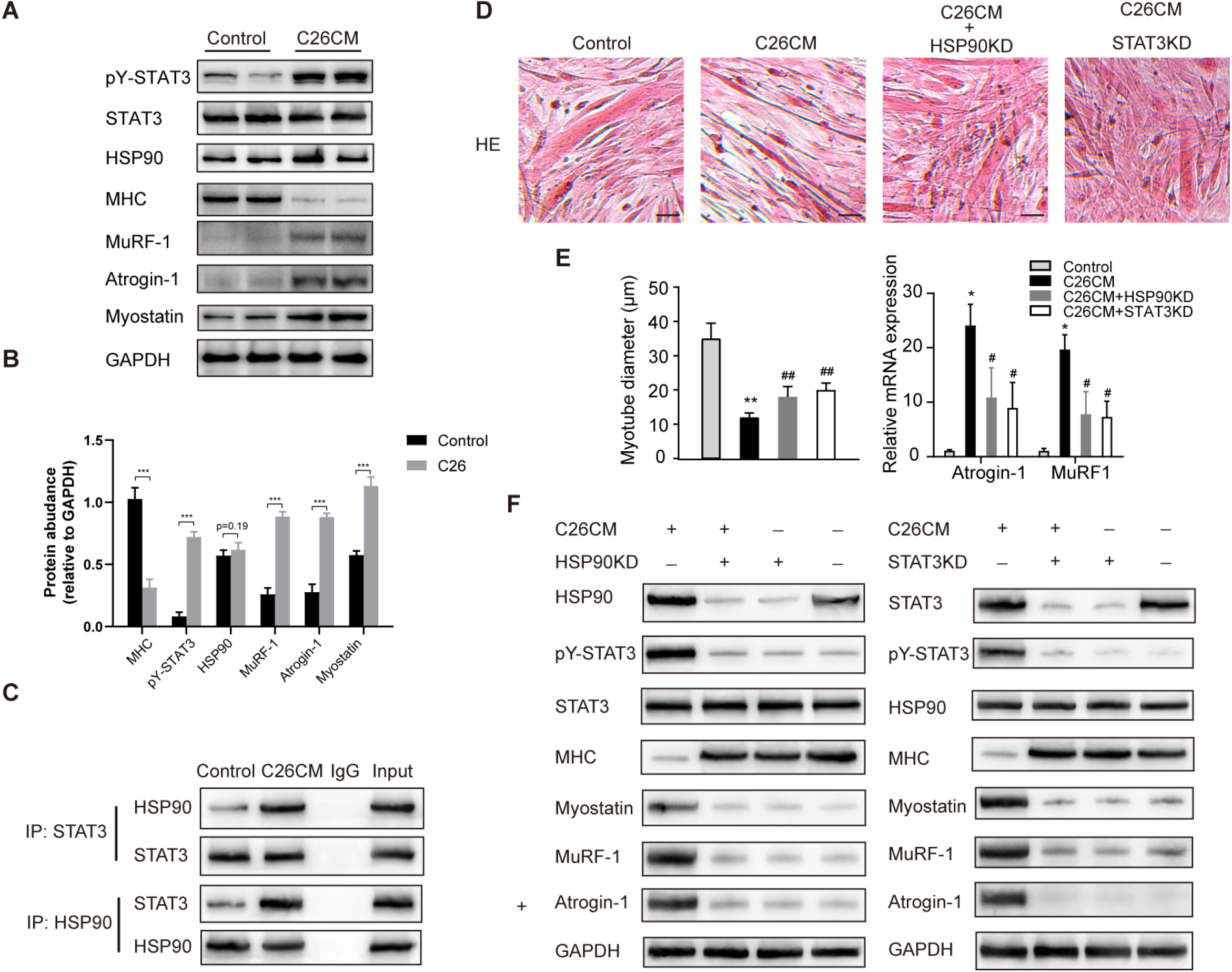
Knocking down of HSP90 or STAT3 prevented C2C12 myotubes from C26 CM-induced atrophy. The expression levels of Atrogin-1, MuRF-1, MHC, HSP90, Myostatin, and phosphorylated STAT3 in the skeletal muscle from C26 CM treated C2C12 myotubes were detected by western blot (A), the bands were quantified by densitometry and normalized to the density of GAPDH (B). The lysates of C2C12 cells treated with C26 CM or DMEM (control) were immunoprecipitated with the STAT3 antibody, the HSP90 antibody, or control rabbit IgG. The immunopellets were detected by immunoblot analysis with the anti-HSP90 antibody and the anti-STAT3 antibody (C). Representative hematoxylin and eosin (HE)-stained C2C12 treated with C26 CM and transfected with the HSP90 siRNA or the STAT3 siRNA (D). The histogram represents the average myotube diameter (μm) (E). qRT-PCR analysis of atrogin-1 and MuRF-1 in the gastrocnemius muscles of mice (F). Atrophy markers in C2C12 were tested by WB analysis when the expression of HSP90 or STAT3 was knocked down by siRNA (G). Data represent the mean ± standard deviation (SD). *p<0.05, **p<0.01 denotes compared to control; #p<0.05, ##p<0.01 denotes compared to C26 CM treated myotubes.

### Disruption of the HSP90-STAT3 interaction by 17DMAG protected C2C12 myotubes from C26 CM-induced atrophy

17DMAG, an inhibitor of HSP90, was then introduced in the C2C12 myotube atrophy model which was established by the treatment of C2C12 myotube with C26 derived CM; the results showed that 17DMAG treatment prevented myotube shrinking, increased the myotube diameter, and enlarged the myotube area (Fig 3A and B, Fig EV2A). Notably, 17DMAG treatment significantly restored the expression of MHC, decreased STAT3 activation, and down-regulated the expression of myostatin, MuRF-1, and Atrogin-1 both at the protein and mRNA levels (Fig 3C-D and Fig EV2D,E). In addition, the colocalization of MHC and myostatin and STAT3 activation were also confirmed by immunofluorescence (Fig 3C). Importantly, 17DMAG disrupted the increase in the binding of HSP90 and JAK2 to STAT3, suggesting that the molecular chaperone activity of HSP90 might prolong the JAK2-STAT3 interaction and cause constitutive STAT3 activation in C26 CM-induced C2C12 cells (Fig 3E). Thus, we concluded that the 17DMAG treatment protected C2C12 myotubes from C26 CM-induced muscle atrophy and catabolism.

**Figure 3.**
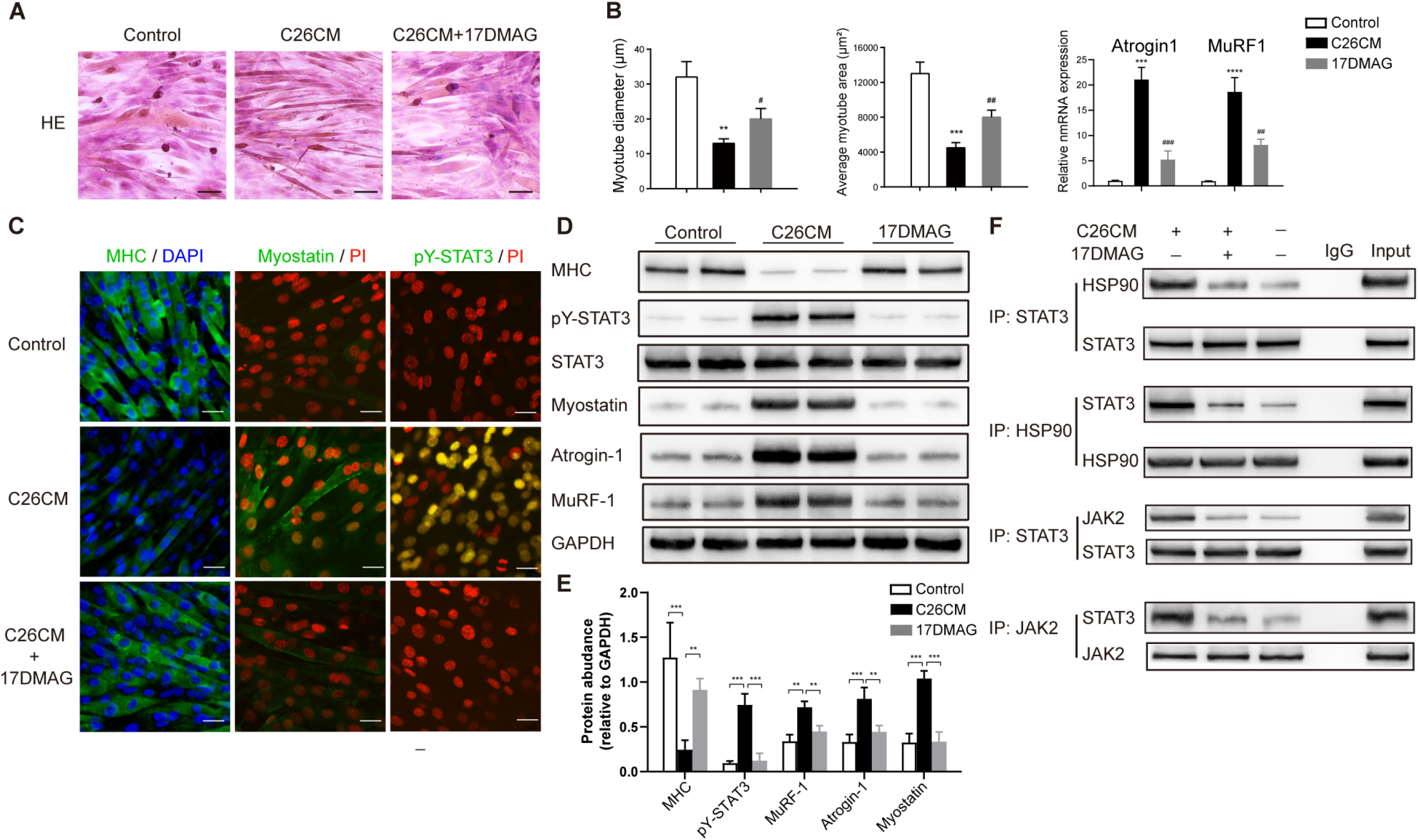
Administration of HSP90 inhibitor 17DMAG prevented C26 CM-induced myotube atrophy and disrupted the binding between HSP90 or JAK2 with STAT3. **A**. C2C12 myotubes were treated with C26 CM, C26 CM plus HSP90 inhibitor 17DMAG (0.1 μmol), or normal media for 24 h. H&E staining shows the morphological changes in C2C12 myotubes. Scale bar: 50 μm. **B**. The histogram represents the average myotube diameter (μm) and myotube area (μm^2^) in control, CM-treated, and 17DMAG plus CM-treated C2C12 myotubes. The number of myotubes was quantified using ImageJ. **C** C2C12 cells were immunostained with the MHC (MF20) antibody. MHC, green, DAPI, blue. The expression and subcellular localization of myostatin and pY-STAT3 in C2C12 myotubes were detected by immunofluorescence staining. Myostatin and pY-STAT3, green, PI, blue. Scale bar: 30 μm. **D** The expression of MHC, pY-STAT3, atrogin-1, and MuRF-1 in control, CM-treated, and 17DMAG plus CM-treated C2C12 myotubes was analyzed by western blot, GAPDH served as a loading control. **E** The lysates of C2C12 cells treated with C26 CM or DMEM (control) were immunoprecipitated with the STAT3 antibody, the HSP90 antibody, or control rabbit IgG. The immunopellets were detected by immunoblot analysis with the anti-HSP90 antibody and the anti-STAT3 antibody. Data represent the mean ± standard deviation (SD). *p<0.05, **p<0.01, ***p<0.001, ****p<0.0001 denotes compared to control; #p<0.05, ##p<0.01, ###p<0.001 denotes compared to C26 CM treated myotubes.

### HSP90 inhibitors ameliorated cachexia in different cachectic mice model

To investigate the protective effect of the HSP90 inhibitor *in vivo*, C26 tumor-bearing cachectic mice were given the HSP90 inhibitor 17DMAG (15 mg/kg, daily) or PU-H71 (50 mg/kg, daily); both drugs were well tolerated by the mice. We observed that the body weights of the mice in the 17DMAG and PU-H71 administration groups were consistently higher than those in the cachectic group (Fig 4A, left). We also observed that 17DMAG or PU-H71 treatment could significantly prolong the overall survival time compared to treatment with the vehicle. Treatment with 17DMAG resulted in a significant shift in the median survival time from 15.5 to 20 days (Fig 4A, right). However, PU-H71 had only a moderate effect on survival time, which was not as significant as that of 17DMAG (Fig 4A, right). 17DMAG or PU-H71 treatment significantly increased the tumor-free body mass compared with the mock control treatment (Fig 4D, left). Notably, neither 17DMAG nor PU-H71 treatment inhibited C26 tumor growth, excluding the possibility that the protective effects of 17DMAG or PU-H71 on body weight were based on its antitumor effect (Fig EV2B). Since 17DMAG showed greater potential for cachexia treatment than PU-H71, it was chosen for further investigation in our work.

**Figure 4.**
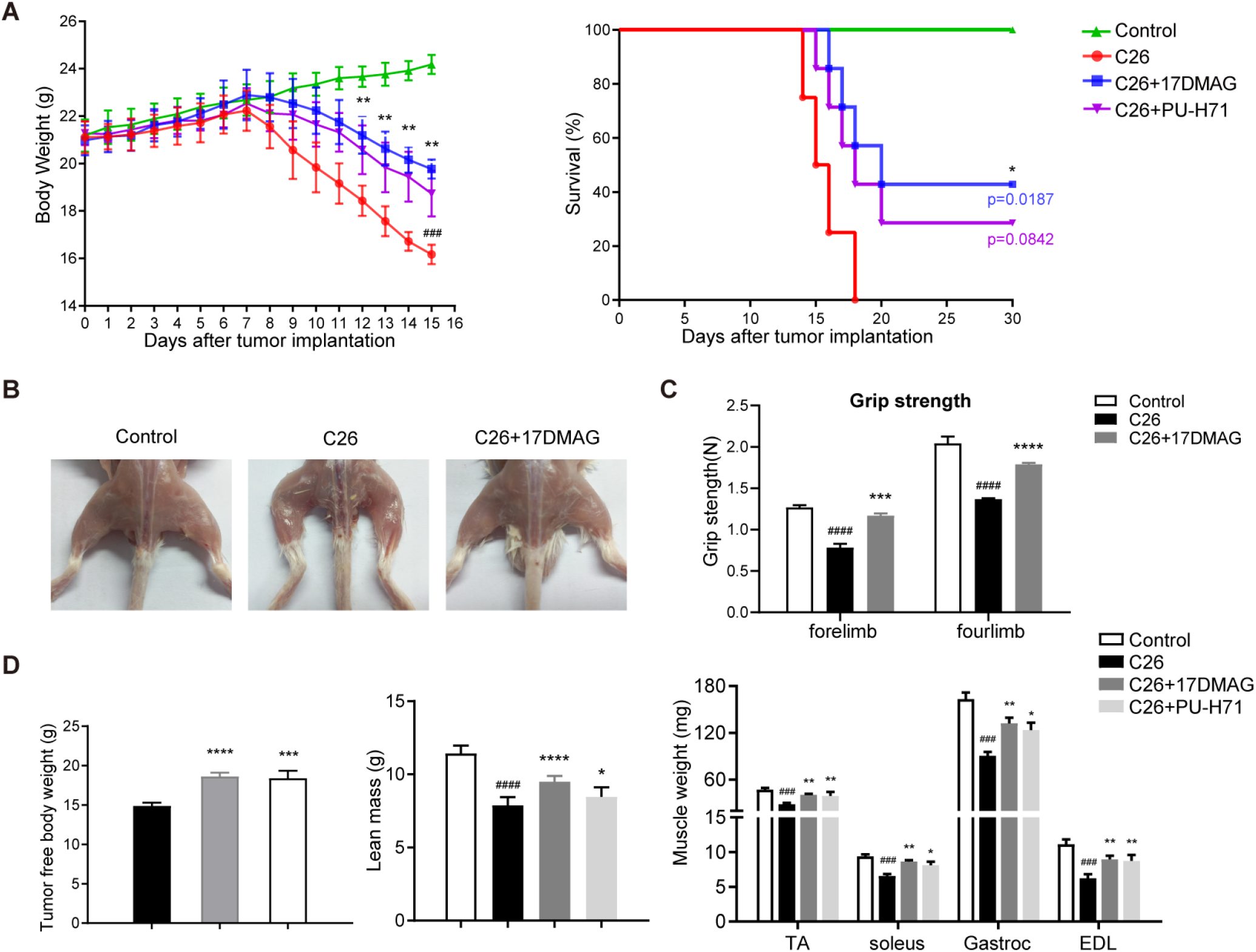
The protective effects of 17DMAG in cachectic muscle wasting in C26 tumor-bearing mice. **A** Left part: Bodyweight changes in normal control mice, C26 tumor-bearing mice, and 17DMAG or PU-H71-treated C26 tumor-bearing mice. Day 0: tumor implantation. Day 7: 17DMAG injection began. Day 15: Terminal sacrifice day. Right part: Survival rates of C26-tumor-bearing mice that received the treatment of 17DMAG or PU-H71 and vehicle control. **B** Representative images of the hindlimbs of normal (left), C26 tumor-bearing (middle), and 17DMAG-treated C26 tumor-bearing (right) mice. **C** Grip strength was measured on day 14. **D** Tumour-free body mass, lean mass weight, and the tibialis anterior (TA), gastrocnemius(Gastroc), soleus, and extensor digitorum longus (EDL) muscles were weighed and compared after dissection with an electronic scale; Data represent the mean ± standard deviation (SD). *p<0.05, **p<0.01, ***p<0.001, ****p<0.0001 denotes 17DMAG vs. C26; #p<0.05, ##p<0.01, ###p<0.001 denotes C26 vs. control.

In C26 tumor-bearing mice, the 17DMAG treatment significantly increased lean body mass and the mass of four major groups of skeletal muscle tibialis anterior (TA), gastrocnemius (Gastroc), soleus, and extensor digitalis longus (EDL), and restored grip strength (Fig 4B-D). In addition, inguinal white adipose tissue (iWAT), which was almost completely absent in C26 tumor-bearing mice (Fig EV2C), was significantly restored by 17DMAG treatment. 17DMAG treatment also resulted in the normalization of the levels of downregulated serum triglycerides (TG) compared with those in the mock treatment vehicle control.

To validate whether 17DMAG has a universal efficiency in different cancer cachexia, the LLC, another Xenograft tumor model of cancer cachexia was applied on C57BL6 mice. As previously reported, the body mass and gastrocnemius muscle mass were decreased in tumor-bearing mice (Gallot *et al*, 2014). 17DMAG administration significantly attenuated the decrease in body mass and muscle mass (Fig EV3C), but final tumor weights of 17DMAG treated tumor-bearing mice were dramatically reduced compared with tumor-bearing control (Fig EV3B), which indicated that 17DMAG not only had a protective effect on cachexia but also inhibited tumor growth in LLC cachectic mice model.

### Inhibition of HSP90 attenuated ubiquitin-proteasome pathway-mediated muscle atrophy in C26 tumor-bearing cachectic mice

Next, we further investigated the protective effect of 17DMAG on cachectic skeletal muscle. H&E staining and morphometric analysis of the gastrocnemius revealed that C26 tumor-bearing mice exhibited an obvious decrease in muscle fiber size, which was markedly blocked by 17DMAG treatment (Fig 5A). The representative frequency distribution of the myofiber CSA showed a rightward shift in 17DMAG-treated mice (the most frequent value for the myofiber CSA was 1500-2000 ìm^2^) vs. C26 tumor-bearing mice (the most frequent value for the myofiber CSA was 1000-1500 ìm^2^) (Fig 5B). The mean CSA of the 17DMAG-treated mice was more than 40% higher than that of C26 tumor-bearing mice (Fig 5C).

**Figure 5.**
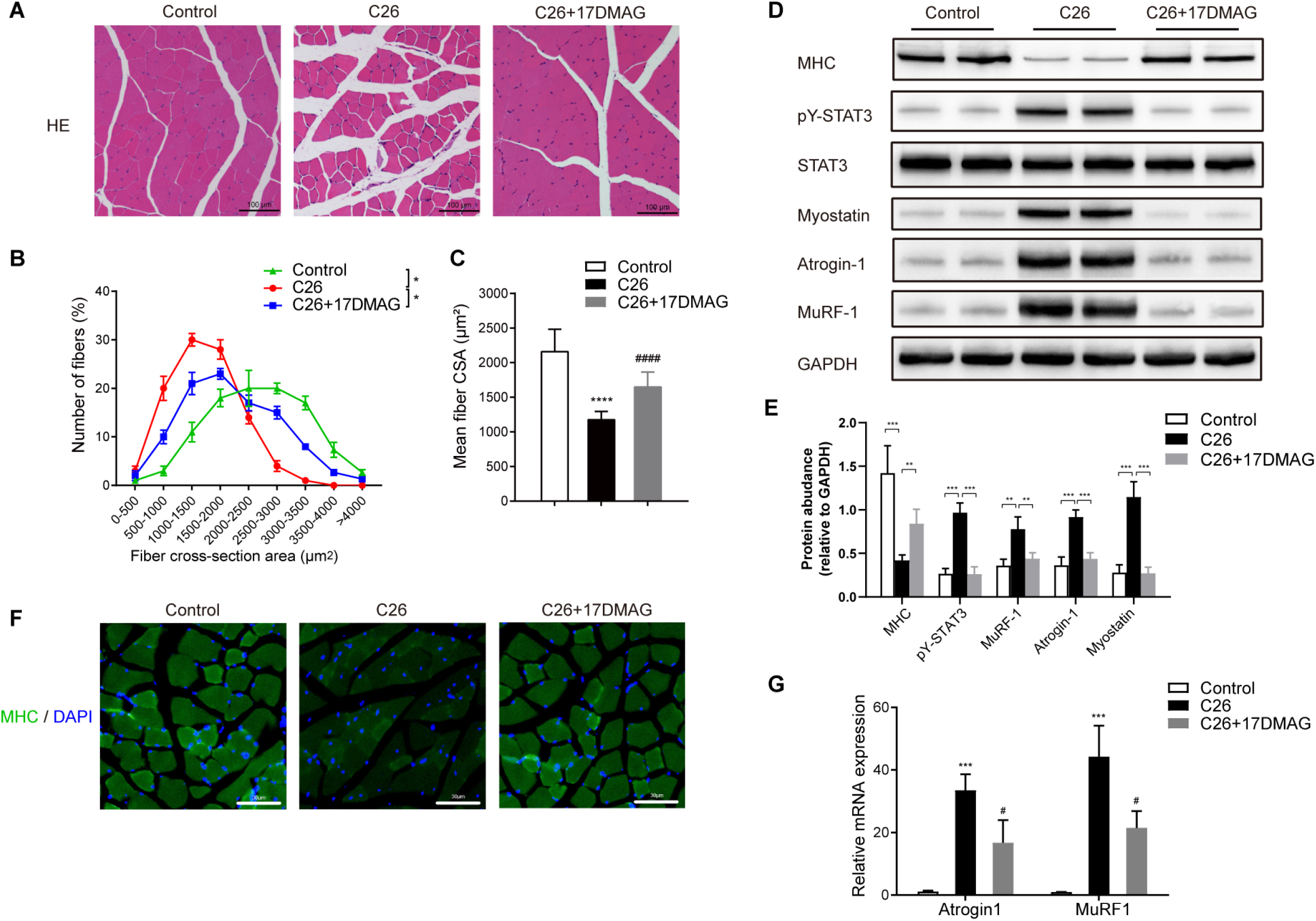
Protective Effects of 17DMAG on fiber cross-sectional area, STAT3 activation, and catabolic pathway suppression. **A** Representative hematoxylin and eosin (H&E) staining showing the morphological changes in the gastrocnemius. Scale bar: 100 μm. **B** Fibre size distribution. **C** Mean fibre cross-sectional area is shown by the χ2 analysis (for CSA); * denotes a significant difference (P < 0.05) compared with normal control; # denotes a significant difference (P < 0.05) compared with C26 mice. **D** 17DMAG treatment altered the expression of myosin heavy chain (MHC), MuRF-1, atrogin-1, and pY-STAT3 in cachectic skeletal muscles from C26 tumor-bearing mice, which were detected by western blot analysis. **E** Gastrocnemius MHC expression was evaluated by immunofluorescence staining. MHC, green, DAPI, blue. Scale bar: 100 μm. **F** Total RNA was extracted from the gastrocnemius from control, 17DMAG-treated, and C26tumor-bearing mice, and the expression levels of MuRF-1 and atrogin-1 in the gastrocnemius were measured by quantitative RT-PCR. Data represent the mean ± SD. ***p<0.001, compared with normal control; #p<0.05, compared with C26 mice.

We also observed that treatment with 17DMAG prevented the striking loss of MHC protein in C26 tumor-bearing mice compared with that in the vehicle control (Fig 5D-E). The significant decreases in STAT3 activation and myostatin, MuRF-1, and atrogin-1 expression in cachectic muscles in C26 tumor-bearing indicated that HSP90 blockade could alleviate muscle wasting by disrupting the STAT3-controlled ubiquitin-proteasome pathway (Fig 5D). These results were further confirmed at the transcriptional level (Fig 5F). Similar results were also observed in PU-H71-treated mice (Fig EV2D).

### STAT3-induced myotube atrophy is FOXO1 dependent

It has been reported that during muscle atrophy, ubiquitin-proteasome pathway activation and the expression of MuRF-1/Atrogin-1 are regulated by the transcription factor FOXO1 (Lokireddy *et al*, 2011; Stitt *et al*, 2004). We, therefore, hypothesized that STAT3 might regulate FOXO1 expression and activation in cachectic skeletal muscle. As shown in Fig. 6a and b, the FOXO1 expression level was significantly increased in cachectic muscle from C26 tumor-bearing mice, and C2C12 myotubes were subjected to C26 CM treatment (Fig 6A and B, FigEV3F). The decreased expression of the inactivated form of pFOXO1 in C26 CM-treated C2C12 myotubes was decreased when STAT3 was knocked down by siRNA (Fig 6B). Next, we transfected a plasmid encoding constitutively activated STAT3 (STAT3-C) (Bromberg *et al*, 1999) into C2C12 myotubes. As expected, STAT3-C transfection significantly upregulated the ubiquitin-proteasome pathway members MuRF-1, Atrogin-1, and myostatin, which was similar to that observed upon C26 CM treatment. Notably, we observed that STAT3-C transfection also significantly increased the level of the activated form of FOXO1 (dephosphorylated) by western blotting (Fig 6C). Immunofluorescence staining confirmed that both STAT3-C and C26 CM were able to increase the nuclear translocation of FOXO1 (Fig 6D).

**Figure 6.**
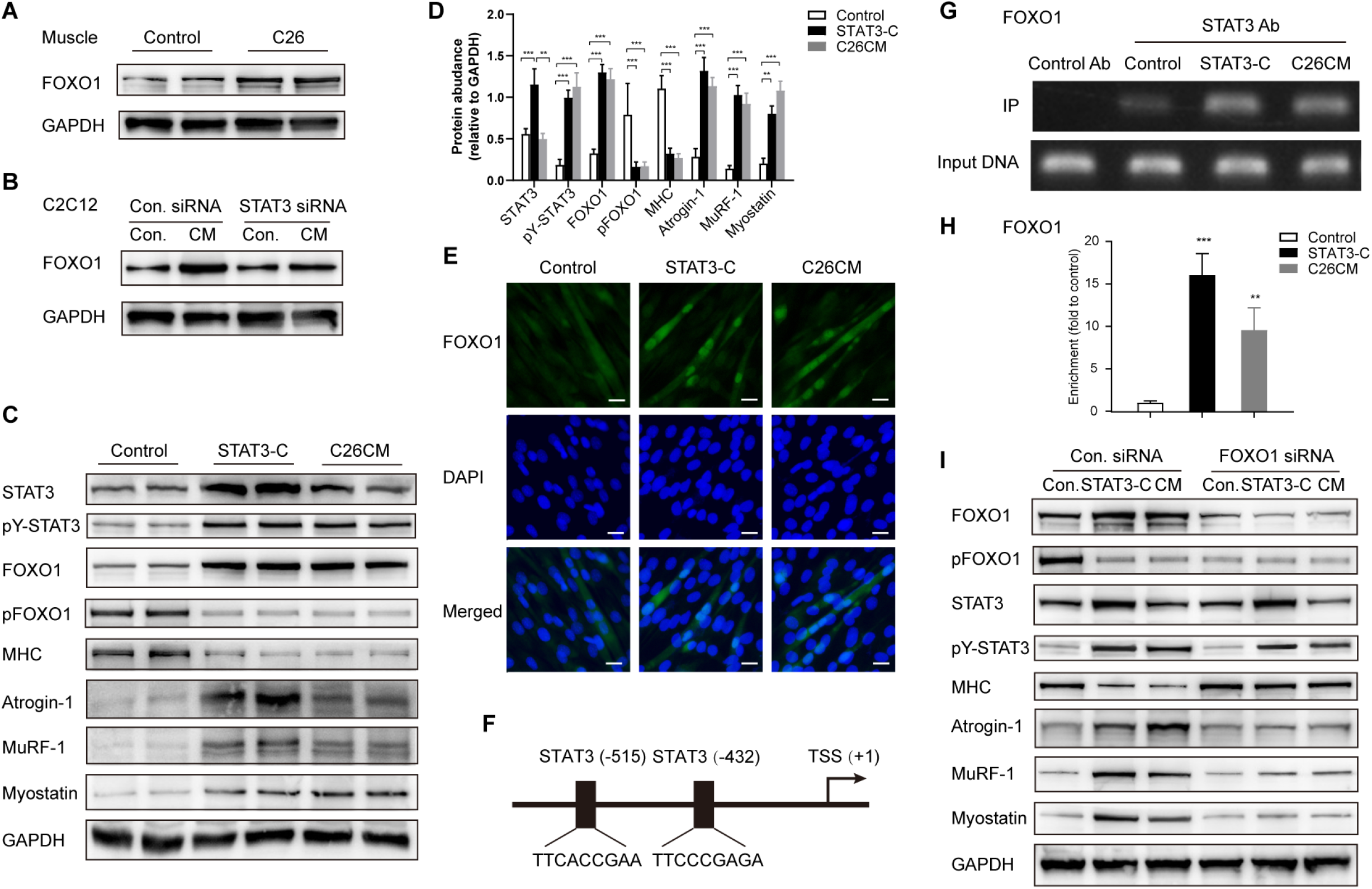
STAT3 regulated FOXO1 expression is responsible for C26 CM-induced myotube atrophy. **A** Measurement of FOXO1 expression of mice muscle and C2C12 myotubes by Western blot. **B** Representative Western blot assay of FOXO1 expression after knocking down of STAT3 by siRNA. **C** Representative Western blot of FOXO1, p-FOXO1, and atrophy genes expression in C2C12 myotube transfected with mutated forms (constitutive activated) of STAT3 overexpression plasmid (Stat3-C). **D** Representative immunofluorescence microscopy images showing the subcellular localization of FOXO1 in C2C12. **E** Schematic structure of the FoxO1 promoter region (NCBI Reference Sequence: NC_000069.5 for FoxO1). Transcription start site (TSS) is denoted by a black arrow. **F-G** Chromatin immunoprecipitation (CHIP) analysis was performed with C2C12, and STAT3 binding to the FoxO1 promoter region was analyzed. Cell lysates were immunoprecipitated with anti-STAT3 Ab or control IgG. Immunoprecipitated and input DNA was analyzed by semi-quantitative PCR (F) or qPCR (G) using primers corresponding to FoxO1 promoter sites. **H** Representative Western blot assay of FOXO1, p-FOXO1, and atrophy genes expression in C2C12 myotube co-transfected with FOXO1 siRNA and STAT3-C recombinant plasmid.

**Figure 7.**
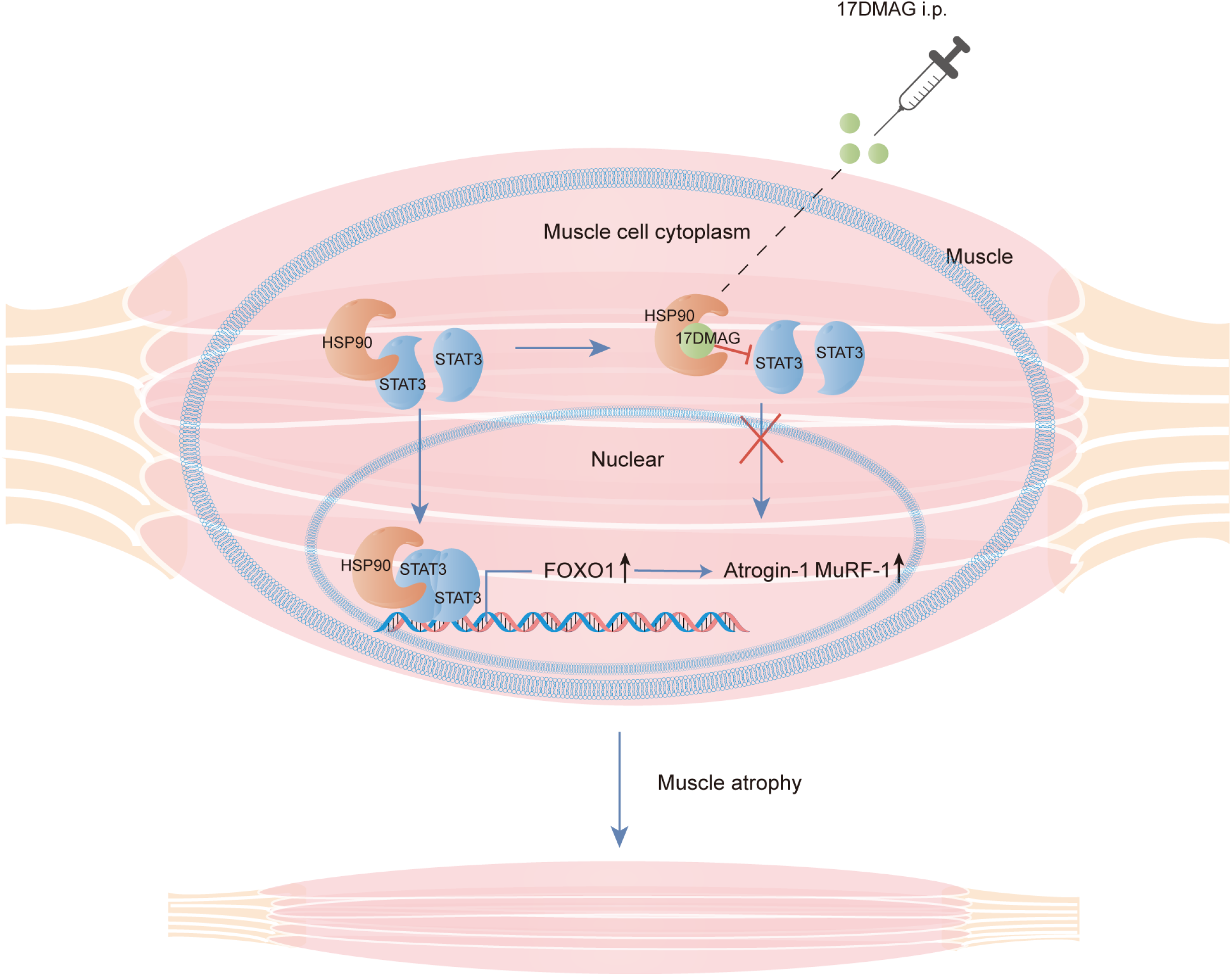
Schematic model of the signaling mechanism of this study.

We next sought to determine whether STAT3 could regulate FOXO1 transcription by direct binding of its promoter. Bioinformatics analysis showed two potential STAT3 DNA binding motifs in the promoter region of the FOXO1 gene (Fig 6E); therefore, chromatin immunoprecipitation assays were performed. As shown in Fig 6F and G, semi-quantitative PCR and qPCR analysis of the DNA immunoprecipitated with the STAT antibody showed the enrichment of the FOXO1 promoter region sequence, indicating direct binding. In contrast, knocking down FOXO1 almost completely abolished STAT3 activation-induced myotube atrophy in both STAT3-C-transfected and C26 CM-treated C2C12 myotube cells (Fig 6H). Our data demonstrated for the first time that in cachectic skeletal muscle, consistent activation of STAT3 could activate the ubiquitin-proteasome pathway in a FOXO1 pathway-dependent manner.

## Discussion

Muscle wasting is one of the hallmarks of cancer cachexia (Penna *et al*, 2018). STAT3 signaling plays crucial role in the regulation of skeletal muscle generation, muscle mass, and repair processes. It is generally believed that transient STAT3 activation in muscles is beneficial to muscle regeneration and hypertrophy, but prolonged STAT3 activation in muscles has been shown to be responsible for muscle wasting (Zimmers *et al*, 2016). Increased STAT3 activation has been observed in skeletal muscles in diverse models of cancer cachexia, including models based on IL-6 administration, C26 colon adenocarcinoma, LLC lung carcinoma, B16 melanoma, and ApcMin/+-induced intestinal cancer (Baltgalvis *et al*, 2008; Bonetto *et al*, 2012; Pretto *et al*, 2015; Puppa *et al*, 2014; Silva *et al*, 2015). However, the clinical association, the factors triggering abnormal STAT3 activation in cachectic muscles as well as the molecular mechanism of muscle atrophy induced by consistent STAT3 activation still need to be delineated.

Most STAT3 molecules exist in the cytoplasm in an unphosphorylated form, and their transcriptional regulatory effects rely on the phosphorylation of the tyrosine 705 residue. The upstream activators of STAT3 include IL-6 family cytokines, growth factors such as EGF, and oncogenes such as the Src family kinases (Yu *et al*, 1995; Zhong *et al*, 1994). Although the excessive release of IL-6 family cytokines, including IL-6, LIF, OSM, and CNTF, has been linked to the pathological development of cancer cachexia (Fearon, 2013). Multiple studies on STAT3 signaling have demonstrated that the stimulation of non-malignant cells with IL-6 family cytokines can trigger only transient STAT3 activation, which is quickly terminated by negative regulators such as PIAS, suppressor of cytokine signaling (SOCS-1, 2, and 3), and cellular phosphatases (SHP-1/2, DUSP22, PRPRD, PTPRT, and PTPN1/2) (Johnson *et al*, 2018; Ram *et al*, 1997; Sengupta *et al*, 1996). Therefore, it is reasonable to speculate that apart from IL-6 family cytokines, other regulatory factors may also be involved in the maintenance of persistent STAT3 activation in cachectic muscles in cancer.

HSP90 is an ATP-dependent molecular chaperone; it has been reported to be one of the most abundant molecules within cells, accounting for nearly 2% of all cellular proteins (Joshi *et al*, 2018). A wide variety of stressful stimuli, including heat shock, UV radiation, and micro-organismal infections, could promote intracellular HSP90 synthesis. During cancer progression, the expression level of HSP90 increases by 2- to 10-fold in tumor cells (Joshi *et al*, 2018). Elevated HSP90 expression has been found in various types of human tumors and is often correlated with increased tumor cell proliferation, poor responses to chemotherapy, and poor survival (Kazarian *et al*, 2017; Ono *et al*, 2018; Whitesell *et al*, 2005; Yun *et al*, 2019). Over 200 oncogenic proteins have been identified as clients of HSP90 (Trepel *et al*, 2010). We and others have demonstrated a direct interaction between STAT3 and HSP90 in multiple cell types, including cancer cells, epithelial cells, fibroblasts, and endothelial cells (Song et al. 2017b; Shah et al. 2002). Here, in this study, we found an increased interaction of HSP90 and STAT3 in skeletal muscle from cachexia patients and experimental mice model. Administration of

HSP90 inhibitors, which blocked the aberrant STAT3 activation, could successfully alleviate cachexia-related muscle wasting phenotype. In addition, we also observed that HSP90 inhibition in cachectic mice could protect against fat loss and increase the number of muscle PAX7+ satellite cells (Fig EV5B). Interestingly, we revealed that the direct interaction of HSP90 with STAT3 could enhance JAK2-mediated STAT3 phosphorylation, thus induce muscle atrophy. The increased HSP90-STAT3 interaction likely cause the prolonged Stat3 activation and muscle atrophy since both HSP90 inhibitor or STAT3 siRNA could successfully alleviate the pathological muscle wasting phenotype both *in-vivo* and *in-vitro*; suggesting that persistent activation of STAT3 rather than other signaling pathway is the major cause of the induction of muscle wasting in cancer cachexia.

An additional goal of this study was to delineate the mechanism by which the prolonged STAT3 activation induces muscle wasting. The pathogenesis of muscle wasting in cancer cachexia has been demonstrated to be the result of an imbalance between the anabolic and catabolic pathways. tThe STAT3 pathway has been found to be a crucial pathway involved in this process. During the catabolic response in skeletal muscles, prolonged STAT3 activation could disturb the sensitivity of the insulin/IGF-1/AKT signaling pathway by enhancing the expression of inhibitory SOCS proteins (Mashili *et al*, 2013; Ueki *et al*, 2004). During the anabolic response, current models indicate that STAT3 could enhance the expression of myostatin via activation of CAAT/enhancer-binding protein (C/EBP) (Zhang *et al*, 2013). Myostatin is a TGF-beta family cytokine that can repress muscle growth and disturb myogenesis. Our study observed the elevated expression of myostatin accompanied by prolonged STAT3 activation, which fully supports our conclusion. However, the most interesting mechanism revealed by our study is that the prolonged STAT3 activation-induced muscle wasting is the result of the overactivation of protein degradation pathways, such as the ubiquitin-proteasome system (UPS). Our data demonstrated that the prolonged activation of STAT3 was correlated with elevated expression of UPS-related muscle-specific E3 ubiquitin ligases, such as atrogin-1 and MuRF1, in skeletal muscle derived from either C26 tumor-bearing cachectic mice or C2C12 myotubes treated with C26 CM. Additionally, the transfection-induced overexpression of constitutively activated STAT3 (STAT3-C) also triggered UPS responses, indicating that constitutive activation of STAT3 itself is enough to induce muscle wasting by activating protein degradation pathways such as the UPS; however, the pathway linking STAT3 and the activation of the protein degradation pathway was not evaluated.

FOXO1, as a key member of the forkhead box protein O (FOXO) transcription factor family, has been found to be widely expressed in various types of metabolism-associated organs, such as the liver and skeletal muscles. In skeletal muscles, the FOXO1 transcription factor represents a master regulator of muscle growth, regeneration, and atrophy. Increased FOXO1 expression and activation has been found in skeletal muscle in energy-deprived states such as fasting, cancer, and severe diabetes (O’Neill *et al*, 2019; Reed *et al*, 2012). Enhanced FOXO1 activation in skeletal muscle was able to trigger muscle atrophy by activation of the UPS and lysosomal proteolysis through the upregulation of FOXO downstream target genes, including atrogin-1 and MuRF1 (Milan *et al*, 2015). Genetic data demonstrated that STAT3 expression and activation are correlated with the intact function of FOXO transcription factors; however, whether FOXO1 is necessary for STAT3-induced muscle wasting or vice versa is still not known. In this study, we demonstrated for the first time that STAT3 could regulate the expression and nuclear translocation of FOXO1 in skeletal muscle, subsequently activating the UPS and triggering muscle wasting in a FOXO1-dependent manner, which could partly explain the pathological mechanism by which consistent STAT3 activation in skeletal muscle could trigger muscle atrophy. Since the bioinformatics study demonstrated that there are several STAT3 consensus motifs in FOXO1 promoter regions, our subsequent ChIP analysis indicated that STAT3 could bind to the FOXO1 promoter and directly regulate FOXO1 gene transcription in skeletal muscles. To test the pathogenic effects of STAT3 on muscle atrophy, a constitutively active recombinant STAT3 mutant expression plasmid, STAT3-C, was adopted in our study. Our results showed that the overexpression of the constitutively activated form of STAT3 in skeletal muscle could mimic the effect of the C26 cell-conditioned medium on inducing the muscle wasting phenotype. Knocking down the expression of FOXO1 in STAT3-C-transfected or C26 cell-conditioned medium-treated myotubes almost completely abolished the activation of the ubiquitin-proteasome system and eliminated muscle wasting, suggesting that STAT3 activation-induced muscle atrophy is indeed FOXO1 dependent.

In summary, our study demonstrated that HSP90 inhibitors could effectively alleviate cancer cachexia-induced skeletal muscle wasting *in vivo*; these effects might be associated with the abnormal activation of the JAK2/STAT3/FOXO1 and ubiquitin-proteasome pathways. The close monitoring of HSP90 levels in cancer patients and the development of strategies to block HSP90 activity could potentially prevent or reverse cachexia development.

## Materials and Methods

### Cell culture

The colon 26 murine adenocarcinoma cell line (C26) and Mouse colon carcinoma (CT26) cells were cultured as a monolayer in RPMI-1640 medium (Life Technology, NY) supplemented with 10% fetal bovine serum, L-glutamine, and 100 units/ml penicillin and streptomycin (HyClone, Logan, UT in a humidified atmosphere containing 5% CO2 at 37 °C. Murine C2C12 myoblasts obtained from the American Type Culture Collection (ATCC) were maintained in Dulbecco’s modified Eagle medium (DMEM) supplemented with 10% fetal bovine serum and antibiotics (100 U/ml penicillin and streptomycin; referred to as growth medium, GM). To induce myotube differentiation, cells were grown to 100% confluence and exposed to DMEM containing 2% horse serum (Life Technology, NY) (referred to as differentiation medium, DM) for up to 4 days.

### Mouse model of cachexia

Six-week-old male BALB/C mice (obtained from the Model Animal Research Center of Nanjing University) were maintained under specific pathogen-free (SPF) conditions and allowed to acclimate for one week. Mice were housed in individually ventilated cages (IVCs) with free access to drinking water and a basal diet under controlled conditions of humidity, light (12 h light/12 h dark cycle), and temperature.

To induce the cachexia mouse model, 100 μl C26 murine adenocarcinoma cells were subcutaneously inoculated into the right flanks of BALB/C mice (5×10^5^ cells/mouse). When the mice lost 20% of their body weight or their tumor volumes reached 1500 mm^3^, the mice were sacrificed. Tumors, muscles, and other organs, such as the liver and hypothalamus, were rapidly dissected, frozen in liquid nitrogen and stored at -80°C until further analyses or fixed with 10% formaldehyde for immunohistological staining. All experiments involving animals were conducted according to protocols approved by the Institutional Animal Care and Use Committee of Nanjing University, in accordance with the guidelines for animal care and use of Jiangsu Province.

### Patients biopsy specimens

The study was approved by the Ethics Committee in the Medical School of Nanjing University (IRB no.: 20200115003). All patients provided written informed consent. Cachexia was defined clinically as documented nonintentional dry weight loss of >5 kg (all >10% of their previous normal weight) at least in ≤3 months (10). All cachectic patients also complained of weight loss; patients’ skeletal biopsy specimens of abdominal muscles were obtained after hepatobiliary surgery. Control samples were acquired from patients undergoing elective orthopedic surgery.

### *In vivo* experiments and tissue collection

To examine the protective effects of the HSP90 inhibitor on cachexia, the cachectic mice received intraperitoneal injection of 17DMAG (HY-12024, MCE,NJ) (15 mg/kg, daily) dissolved in 1% DMSO + 30% PEG300 + 1% Tween 80 or PU-H71 (HY11038,MCE,NJ) (50 mg/kg, daily) dissolved in PBS (as the solvent control) beginning on day 7 when the tumor became palpable. For the survival experiment, thirty mice were randomly divided into four groups (normal control, C26, C26 + 17DMAG, and PU-H71 groups), and each group contained 10 mice. The mice were followed until the endpoint criterion was fulfilled or at least 30 days had elapsed after the inoculation of C26 cells to determine the survival rate.

The endpoint criterion combined the detected weight loss with the overall condition of the mouse. To evaluate the overall health status of the mice, the following aspects were considered in addition to weight loss: appearance, posture, and natural and provoked behavior (inactivity, dyskinesia, and reduced response to external stimuli). Mice were euthanized when two researchers confirmed the fulfillment of the endpoint criteria.

To study the potential mechanisms involved, another experiment was conducted with a predetermined endpoint at day 15 after C26 cell inoculation. The numbers of mice and groups were the same as those used in the survival experiment. The following data were collected: body weight, food intake, tumor weight, grip strength, and survival rate. Once the tumors were removed, the gastrocnemius was dissected, harvested, weighed, and then fixed in 4% paraformaldehyde (Sigma-Aldrich, St. Louis, MO) for subsequent histological analysis.

### Assessment of grip strength

Skeletal muscular strength in mice was quantified with the grip strength test. At the end of the experiment, the forelimb grip strength of each of the four limbs was measured with a Digital Grip Strength Metre (Nanjing Biomedical Research Institute of Nanjing University). Mice were held by the tail and allowed to grasp a grid with their forepaws or with all four paws. The mice were gently pulled by the tail until they released their grip. Five measurements at 3 min intervals were recorded, and the average value for each mouse was determined.

### Histological analysis and immunofluorescence

Procedures for the determination of the fiber cross-sectional area (CSA) were conducted as previously described (Park *et al*, 2013). Briefly, fixed transverse sections (7 μm) were cut from the mid-belly of the gastrocnemius with a cryostat at -20°C and then stained with hematoxylin and eosin (H&E). Images were obtained at 40× magnification and quantified using ImageJ software (National Institutes of Health, Bethesda, MD). The CSA and number of myofibers with central nuclei among the individual myofibers were determined by using ImageJ 1.48 software in five random fields for each section.

For immunofluorescence analysis, cells were seeded onto sterile preprocessed glass coverslips that were precoated with 1% gelatin. After C2C12 myoblasts had differentiated into myotubes, the cells were washed twice with PBS, followed by fixation in 4% paraformaldehyde for 15 min. After being rehydrated in PBS, the cells were blocked for 30 min in 1% bovine serum albumin (BSA) in PBS containing 0.2% Triton-X (PBST). Then, the cells were incubated with an anti-myosin heavy chain (MHC) (#MAB4470, R&D Systems,MN) (1:100) in 1% BSA/PBST overnight at 4°C. Cells were then incubated with a fluorescence-labeled secondary anti-mouse antibody (1:1000) and DAPI (1:1000) at room temperature for 1 h. The specimens were examined under an FV10i laser scanning confocal microscope (Olympus, Tokyo, Japan).

### Western blot and immunoprecipitation assays

Tissues and cell proteins were extracted in ice-cold RIPA lysis buffer consisting of 150 mM NaCl, 50 mM Tris-HCl, 0.5% sodium deoxycholate, 200 mM NaF, 200 mM PMSF, 1.0% NP40, 1 mM EDTA, and protease and phosphatase inhibitor cocktail (Life Technology, NY). The protein concentration in the supernatants was determined with a BCA Kit (Pierce, Rockford, IL). Then, 20 µg of total protein was separated by electrophoresis in a 10% SDS polyacrylamide gel and transferred to a PVDF membrane (Millipore, Billerica, MA) for western blot analysis.

The signals were detected with the following antibodies by following standard procedures: STAT3 (#9145,CST, MA), phosphorylated STAT3 (#9139,CST, MA), HSP90 (ab13492, Abcam,MA), myostatin (ab71808, Abcam,MA), CD63 (ab8219,Abcam, MA), atrogin-1 (sc166806,Santa Cruz, MA), MuRF-1 (sc398608,Santa Cruz, TX), MHC (#MAB4470, R&D Systems,MN), GAPDH (ab181602,Abcam, MA).

For the immunoprecipitation assays, cells were lysed in WB/IP buffer, and proteins were immunoprecipitated with the indicated antibodies. Precleared Protein A/G Plus-Agarose beads (Life Technology, New York, NY) were incubated with the immunocomplexes overnight and washed three times with lysis buffer. The immunoprecipitates were subjected to SDS-PAGE followed by immunoblotting.

### Transfection of small interfering RNAs (siRNAs) and overexpression plasmids

siRNA transfection was performed using Lipofectamine RNAiMAX transfection reagent (Life Technology, New York, NY) by following the manufacturer’s guidelines. Briefly, C2C12 cells were seeded into a six-well plate. After four days of differentiation, 5 pmol of siRNA was diluted with Opti-MEM and mixed with the transfection reagent, and the mixture was added to each well. After transfection for 24 h, the cells were challenged with different treatments; some were co-cultured with C26 cell-conditioned media (C26 CM), and some were cultured in DMEM. The siRNAs specific to HSP90aa1 were purchased from Thermo (#159050,Life Technology, New York, NY). The siRNAs specific to FOXO1 were purchased from Santa Cruz (#56458,Santa Cruz, TX).

The constitutively activated STAT3 plasmids (STAT3-C Flag pRc/CMV) were purchased from Addgene (Plasmid #8722). C2C12 cells were transiently transfected with STAT3-C plasmids using the Lipofectamine™ 2000 Transfection Reagent according to the manufacturer’s instructions (Life Technology, New York, NY).

### Chromatin immunoprecipitation (ChIP) analysis

The ChIP assay was performed according to the manufacturer’s instructions (P2078,Beyotime,China). C2C12 cells were fixed with 1% formaldehyde at 37°C for 10 min, lysed, and sonicated. Soluble chromatin was coimmunoprecipitated with anti-STAT3 antibodies or an equal amount of rabbit IgG. The immunoprecipitated and input DNA were subjected to PCR and qPCR using FOXO1 promoter site-specific primers (5′-CGACTTCAACACCTCATCGCTTC-3′ and 5′-AGGCGCGCAGATCCTTCGGTGA-3′). Q-PCR was performed with Power SYBR Green PCR Master Mix (Life Technology, New York, NY). The fold change in enrichment relative to the input DNA was calculated using the comparative Ct method (ΔΔCt).

### Real-time PCR assay

Total RNA from either snap-frozen muscles or cultured cells was isolated using an RNeasy Micro Kit (Qiagen, Hilden, Germany). The mRNA was reverse-transcribed into cDNA with PrimeScript RT Master Mix (TaKaRa, Otsu, Japan) and subjected to quantitative real-time PCR with SYBR Green PCR Master Mix (Life Technology,New York,NY). The relative expression of the target genes was normalized to that of GAPDH and calculated with the ΔΔCt method. The following primers were used: MuRF-1: forward: 5′-ACCTGCTGGTGGAAAACATC-3′, reverse: 5′-AGGAGCAAGTAGGCACCTCA-3′; Atrogin-1: forward: 5′-ATTCTACACTGGCAGCAGCA-3′, reverse: 5′-TCAGCCTCTGCATGATGTTC-3′; GAPDH: forward: 5′-ACCCTTAAGAGGGATGCTGC-3′, reverse: 5′-CGGGACGAGGAAACACTCTC-3′.

### Clinical data mining

Data from The Cancer Genome Atlas (TCGA), including gene expression data and patient survival data, were downloaded from the TCGA web server with GDC Client software. For HSP90AA1 expression analysis, cancer patient data were obtained from TCGA, and gene expression data for normal tissue were obtained from the Genome Tissue Expression (GTEx) portal.

### Statistical analysis

The tissue weights of different groups were compared using Student’s t-test. Longitudinal body weight or muscle mass differences were analyzed by repeated measurement analysis of variance (ANOVA). The survival rate difference was determined with the chi-square test. P values < 0.05 were considered statistically significant.

## Author contributions

M.N, S.S, and Z.S designed and performed the experiments, analyzed data, and drafted the manuscript. W. P and C.Z. performed a part of the animal experiments. L.W, L.L, and Y.D performed a part of the *in vitro* experiments. W.C, Q. G, and H.W conceived the projects, designed the experiments, analyzed data, and wrote the manuscript. All authors read and approved the final manuscript.

## Acknowledgments

This work was supported by grants from the National Natural Science Foundation of China (Nos. 82070912, 81773326 and 81471095), the Key Project of Research and Development of Ningxia Hui Autonomous Region of China (No. 2017BN04), a grant from the Natural Science Foundation of Jiangsu Province China (No. BK20171347 and BE2019676) and the Qing Lan Project of Jiangsu Province China. We thank Prof Erguang Li for his critical review and thoughtful discussion of the manuscript.

## Expanded View Figure legends

**Figure EV1.**
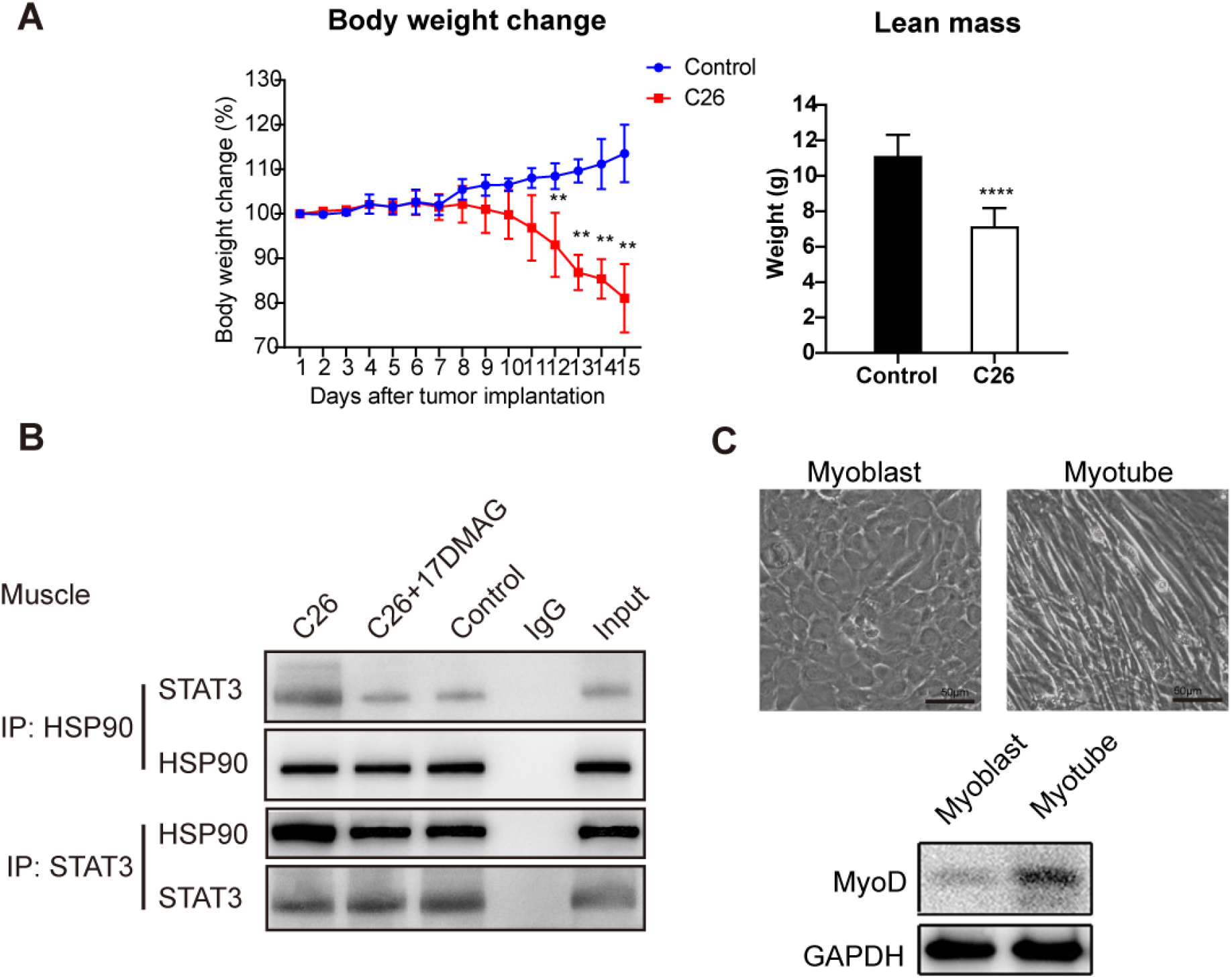
**A**. Body weight change of C26 tumor-bearing mice compared with non-tumor bearing control. **B**. Co-immunoprecipitation assay of HSP90 and STAT3 in skeletal muscles from normal and C26 cachectic mice with or without 17DMAG treatment. **C**. Phase-contrast images of C2C12 myoblasts and myotubes; Detection of MyoD expression in myoblasts and myotubes by western blot analysis.

**Figure EV2.**
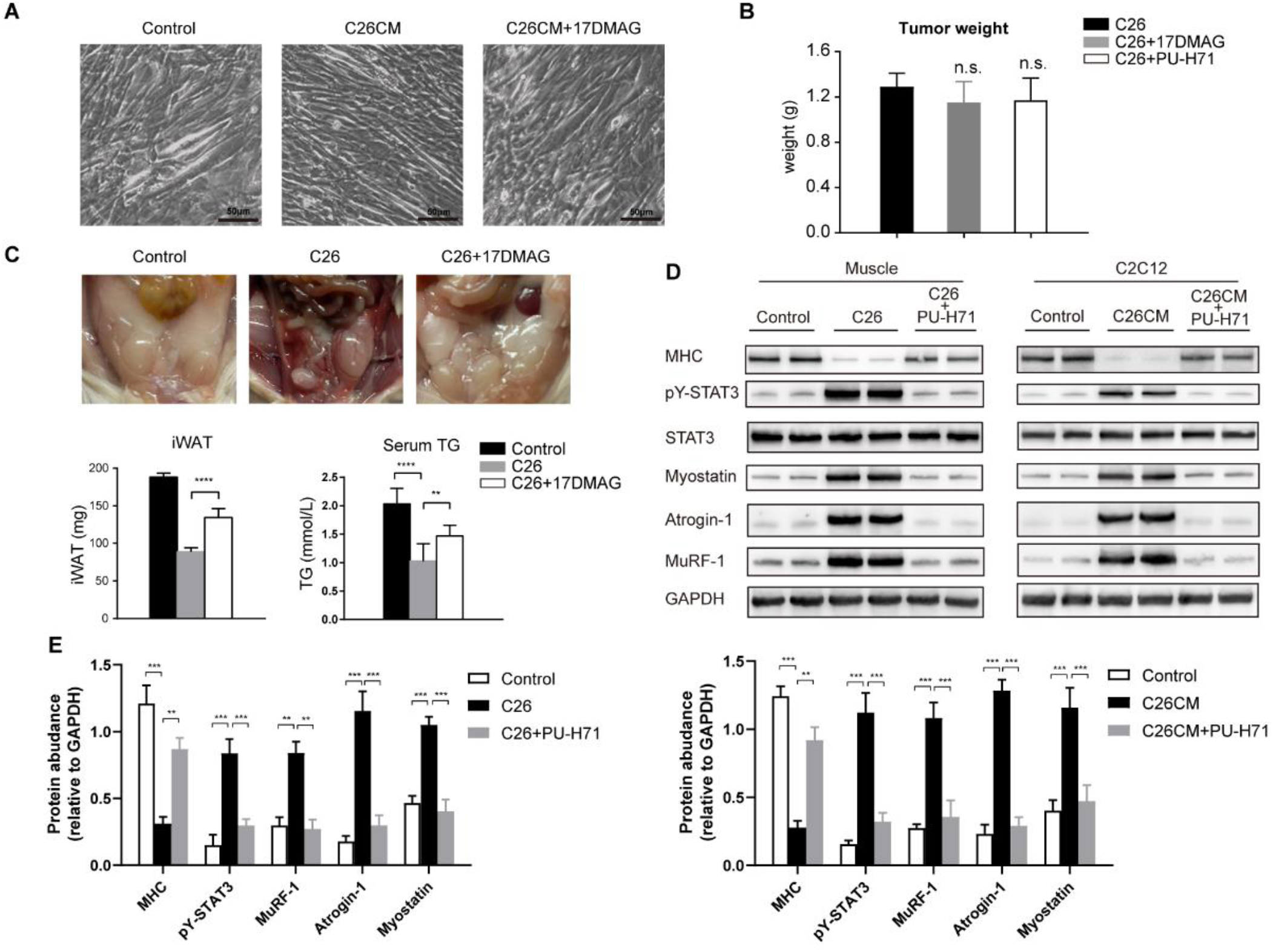
**A**. Phase-contrast images of C2C12 cells treated with C26 CM or 17DMAG and the non-CM-treated control. **B**. Final tumor weights of 17DMAG-treated and C26 tumor-bearing mice were measured at necropsy. m**C**. Inguinal white adipose tissue (iWAT); iWAT weight and serum TG level. **D**. Representative western blot of MHC, pY-STAT3, atrogin-1/MAFbx and MuRF-1 in muscle and C2C12 treated by PU-H71. Data represent the mean ± SD. **p<0.01; ***p<0.001; ****p<0.0001.

**Figure EV3.**
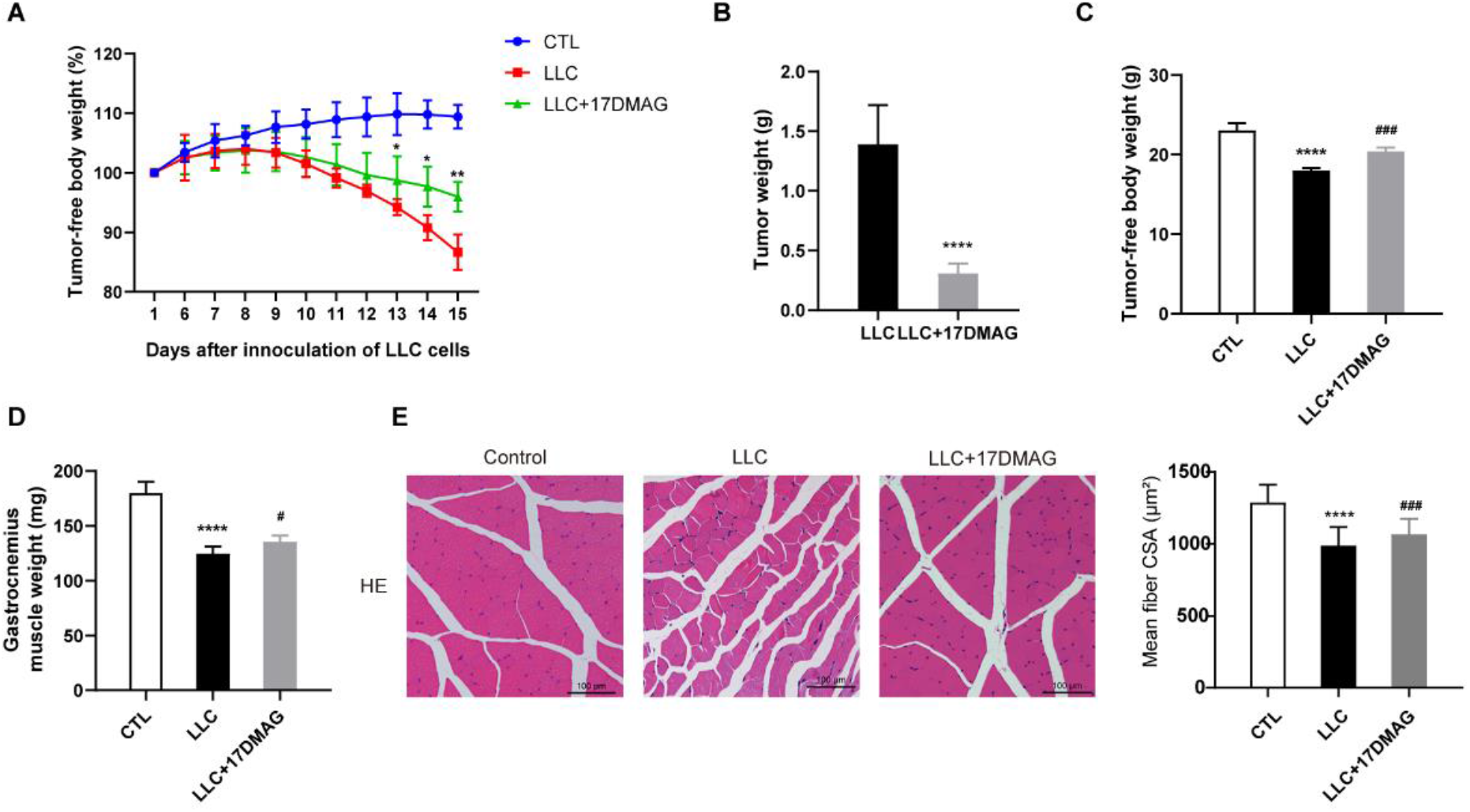
**A**. Time course change of tumor free body weights of 17DMAG-treated and LLC tumor-bearing mice. **B**. Final tumor weights of 17DMAG-treated and LLC tumor-bearing mice were measured at necropsy. **C**. Tumor free body weights of 17DMAG-treated and LLC tumor-bearing mice. **D**. Weight change of gastrocnemius Muscle 17DMAG-treated and LLC tumor-bearing mice. **E**. Representative H&E staining showing the morphological changes in the gastrocnemius Muscle. Scale bar: 100 μm. **F**. Transcription level of FOXO1 in C26 mice muscle was detected by qPCR. **G**. PAX7 immunofluorescence revealed the activation of satellite cells in skeletal muscle from the mouse groups described above. PAX7: green; DAPI: blue; scale bar: 30 μm. Data represent the mean ± SD. **p<0.01

